# The Future is Less Concrete than the Present: A Neural Signature of the Concreteness of Prospective Thought Is Modulated by Temporal Proximity during Intertemporal Decision-Making

**DOI:** 10.1101/2021.02.13.431095

**Authors:** Sangil Lee, Trishala Parthasarathi, Nicole Cooper, Gal Zauberman, Caryn Lerman, Joseph W. Kable

## Abstract

Why do people discount future rewards? Multiple theories in psychology argue that future events are imagined less concretely than immediate events, thereby diminishing their perceived value. Here we provide neuroscientific evidence for this proposal. First, we construct a neural signature of the concreteness of prospective thought, using an fMRI dataset where the concreteness of imagined future events is orthogonal to their valence by design. Then, we apply this neural signature in two additional fMRI datasets, each using a different delay discounting task, to show that neural measures of concreteness decline as rewards are delayed farther into the future.

**Significance Statement:** People tend to devalue, or discount, outcomes in the future relative to those that are more immediate. This tendency is evident in people’s difficulty in making healthy food choices or saving money for retirement. Several psychological theories propose that discounting occurs because delayed outcomes are perceived less concretely that more immediate ones. Here we build a brain decoder for the concreteness of future thought and use this unobtrusive measure to show that outcomes are processed less concretely as they occur farther into the future.

---

Many of the most important choices we make in our daily lives involve tradeoffs between the present and future. Should I spend money now or to save it for retirement? Can I forego the pleasure of eating this dessert now in order to reach my weight loss goal and improve my health? In such intertemporal decisions, humans tend to devalue, or discount, outcomes in the future; a phenomenon known as delay discounting. In the laboratory, this tendency can be measured by presenting participants with choices between a smaller monetary amount available immediately or a larger monetary amount available after a delay. Patience as measured by laboratory intertemporal choice tasks predicts other important aspects of life such as drug and alcohol abuse, educational attainment, and personal finances (Alessi & Petry, 2003; Anderson & Mellor, 2008; Brañas-Garza, Georgantzís, & Guillén, 2007; Kirby, Petry, Nancy, & Bickel, Warren, 1999; Krain et al., 2008; Lejuez, Aklin, Bornovalova, & Moolchan, 2005; Lejuez et al., 2003; Schepis, McFetridge, Chaplin, Sinha, & Krishnan-Sarin, 2011; Shamosh & Gray, 2008; Urminsky & Zauberman, 2015).

Why, however, are delayed outcomes fundamentally less desirable? Psychologists have long pondered this important question. Several theories suggest that one potential explanation is that future outcomes are less concrete. Rick and Lowenstein (2008) have pointed out that in many intertemporal decisions, delayed outcomes are intrinsically less tangible than sooner ones. For example, while a calorie-rich dessert yields immediately perceivable pleasure for the eater, the promise of better future health is less appreciable. Similarly, construal level theory proposes that even when future outcomes are not intrinsically less tangible, people tend to use a process of high-level construal when thinking about future events that leads to them being represented in a relatively more abstract way (Liberman & Trope, 2014; Trope & Liberman, 2010). In contrast, when people consider sooner events, they use low-level construal and represent them in a relatively more concrete manner. Many behavioral studies have provided some support for the central claim that the same outcome is perceived less concretely when it occurs farther in the future rather than more immediately (for review, see Liberman & Trope, 2014), and linked these to changes in representation of delay discounting (Malkoc & Zauberman, 2006; Malkoc, Zauberman, & Bettman, 2010). However, an ideal test of these construal effects in discounting would measure concreteness on-line and non-obtrusively, while people are making intertemporal decisions.

Functional brain imaging has the potential to provide such a non-obtrusive online test of whether future outcomes are perceived as less concrete during intertemporal decision-making. Yet, while many fMRI studies have compared brain activity for sooner versus later outcomes (for review, see Carter, Meyer, & Huettel, 2010), attributing any neural differences specifically to concreteness requires ruling out other potential sources for these differences. Perhaps the most important and obvious difference between sooner and later outcomes that could drive neural activity is that sooner outcomes are valued more highly than delayed ones; that is, brain activity selectively responding for sooner versus later outcomes may reflect valuation, not necessarily concreteness. Indeed several previous studies that have compared sooner and later outcomes have found increased activity in the medial prefrontal cortex (mPFC) and posterior cingulate cortex (PCC), two regions with well-established roles in valuation (Bartra, McGuire, & Kable, 2013; Cooper, Kable, Kim, & Zauberman, 2013; Lee, Parthasarathi, & Kable, 2020; Mitchell, Schirmer, Ames, & Gilbert, 2011; Tamir & Mitchell, 2011). Instead, what is required is a neural signature that is specifically predictive of concrete versus abstract prospective thought, and independent of positive versus negative evaluation.

In the current study, we use a recently developed novel method (Lee, Bradlow, & Kable, 2021) to construct a whole-brain multivariate neural predictor of the concreteness of imagined future events. To train this predictor, we use a prospective imagination dataset (Lee et al., 2020), in which the concreteness (high versus low) and valence (positive versus negative) of imagined future events were orthogonal, such that the subsequent neural predictor was specific to concreteness and not valence. We then applied the whole-brain neural predictor of concreteness in two separate delay discounting task datasets with different evaluation schemes (bidding vs. choice) to test whether the temporal distance of monetary options in intertemporal decision-making modulates the neural signature of concrete versus abstract imagination.

## Methods

### Prospection Dataset

We used a dataset from (Lee et al., 2020) to develop a whole brain predictor of the concreteness of imagined future events. This study examined neural activity associated with the valence (positive versus negative) and concreteness (high versus low) of imagined future events. Twenty-four participants underwent fMRI scanning while imagining thirty-two different future scenarios. In a 2×2 design (positive versus negative valence crossed with high versus low concreteness), eight different unique scenarios were selected for each condition based on pilot testing. Each scenario was repeated twice during the experiment. Participants completed four runs and imagined sixteen scenarios per run. Each trial involved up to 5 seconds of participants reading the scenario cue, 12 seconds of imagination, and up to 14 seconds in which participants rated the concreteness and valence of the imagined event on a 7-point Likert Scale (7 seconds each). The trial duration was buffered such that the time the participants did not use in the cue phase and the rating phase was appended to the ITI at the end of the trial to make the total duration of a single trial 34 seconds.

### Delay Discounting Datasets

We applied the neural predictor of concreteness developed in the prospection dataset in two different delay discounting datasets to test whether the neural signature of concreteness is modulated by delay during intertemporal decisions. We use one bidding dataset and one choice dataset to evaluate the robustness of the results to different task structures. The first dataset we used was from Cooper et al., 2013, which involved bidding on delayed rewards. A total of forty participants were asked on each trial to indicate an immediate monetary amount that they would feel was equivalent to receiving $75 after a given delay, varying from 14 to 364 days. Each trial began with a screen of the form “I feel indifferent between receiving $75 in 28 days and receiving ____ now”. After the prompt was shown for 3 to 5 seconds, participants were then allowed a maximum of 10 seconds to use a button pad to indicate their immediate equivalent amount within a range of $0 to $75. Each participant went through four scan runs, each of which involved twenty-six questions at different delays, ranging from 14 to 364 days. We removed one participant who bid $75 for all trials regardless of delay, as we were not sure whether the participant understood the task. An advantage of this dataset is that it presents participants with the exact same reward amount at varying delays, thereby allowing us to test whether the neural signature of imagination concreteness is modulated by the delay. The flipside of this advantage is that only the delay, and not the amount, of the delayed reward is varied across trials. This limitation is addressed in the second dataset below.

The second dataset we used was from Kable et al. 2017, which investigated the effects of cognitive training on neural activity during economic decision-making. Here we use the data from the intertemporal choice task in the first, baseline, scanning session. One hundred sixty-six participants completed four runs of the intertemporal choice task while being scanned. Each run consisted of thirty binary choices between a smaller immediate reward of $20 today that was held constant throughout the entire session and a larger delayed reward (e.g., $30 in 7 days) that varied in amount and delay from trial to trial. On each trial, the delayed option was shown on the screen; the immediate option was not displayed. Participants pressed the left/right buttons on a button pad to indicate whether they would like to accept the delayed option shown on the screen and forego the immediate reward of $20, or to reject the delayed option and take the immediate reward of $20. Participants had up to 4 seconds to respond, and after their response, a checkmark was shown on the screen if they accepted the delayed reward and an X was shown on the screen if they rejected it.

### Image acquisition

For all datasets, the images were collected with a Siemens 3T Trio scanner with a 32-channel head coil. High-resolution T1-weighted anatomical images were acquired using an MPRAGE sequence (T1 = 1100ms; 160 axial slices, 0.9375 × 0.9375 × 1.000 mm; 192 × 256 matrix). T2*-weighted functional images were acquired using an EPI sequence with 3mm isotropic voxels, 64 × 64 matrix, TR = 3,000ms, TE = 25ms (TE = 30ms for Cooper et al., 2013). The prospection dataset’s EPI sequence involved 44 axial slices with 181 volumes (Lee et al., 2020), the intertemporal bidding dataset’s EPI sequence involved 45 axial slices with 150-152 volumes (Cooper et al. 2013), and the intertemporal choice dataset’s EPI sequence involved 53 axial slices with 104 volumes (Kable et al. 2017). Lee et al. (2020) and Kable et al. (2017) collected B0 fieldmap images for distortion correction (TR = 1000ms, TE = 2.69 and 5.27ms for prospection dataset and TR = 1270ms, TE = 5 and 7.46ms for the intertemporal choice dataset).

### Image preprocessing

All datasets were preprocessed via fMRIPrep (Esteban et al., 2019). The details on the preprocessing pipeline, as generated by fMRIPrep and unaltered, are available in the supplemental materials. In short, all BOLD runs were motion-corrected, slice-time corrected, b0-map unwarped, registered and resampled to a MNI 2mm template. fMRIPrep does not perform smoothing, so it was manually performed after estimating single trial activities (see below).

### BOLD deconvolution

We used beta-series regression (Rissman, Gazzaley, & D’Esposito, 2004) to estimate the BOLD activity associated with each trial in each of the three datasets. In the prospection dataset, we estimated the BOLD activity during the imagination period of 12 seconds. The regressors were time-locked to imagination time onset with an event duration of 12 seconds and convolved with a double gamma HRF function. In the intertemporal bidding dataset from Cooper et al. (2013), the regressors were time-locked to the onset of the response period (when participants can input their bids) with event duration of 0.1 seconds and convolved with a double gamma HRF function. Finally, in the intertemporal choice dataset from Kable et al. (2017), the regressors were time-locked to the trial onset period with event duration of 0.1 seconds and convolved with a double gamma HRF function. In this dataset only, the last trial of each run was excluded from analysis because the BOLD activity of the last trial was often not observed due to the termination of the scan. After the single trial coefficients were estimated, all images were smoothed with a FWHM 8mm gaussian filter, which is the standard smoothing kernel for SPM.

### Concreteness prediction map

To create a whole brain predictor of the concreteness of imagined future events, we used thresholded partial least squares (T-PLS; Lee et al., 2021). T-PLS is similar in approach to other methods for constructing whole predictors that use principal components analysis (PCA) to reduce the dimensionality of the data followed by regression (Chang, Gianaros, Manuck, Krishnan, & Wager, 2015; Wager, Atlas, Leotti, & Rilling, 2011; Wager et al., 2013). The key advantage of T-PLS over PCA-based methods is that partial least squares (PLS) is used for data-reduction. PLS components maximally explain the covariance between the predictors and the outcome, whereas PCA components only explain the variance of the predictors. Thus, PLS yields data-reduction that is more pertinent to prediction.

We built the whole-brain predictor of concreteness in three steps (**Fig. 1**). First, we performed PLS to extract components that maximally explain the covariance between the single trial images and the binary concreteness trial categories (high versus low). These components consist of a map of weights for each voxel in the brain. PLS also automatically yields coefficients for each component that are equivalent to the regression coefficients one would obtain from regressing the dependent variable on the components. We also calculated the t-statistics of each component as one would get from a regression model (here, given the large number of observations, we assume that the t-statistics are approximations of z-statistics). In the second step, we back-project the PLS coefficients and z-statistics into the original voxel space by multiplying them with the PLS weight maps. This yields coefficients for each of the brain voxels for easier interpretation. In the final step, we used the back-projected z-statistics of each voxel to rank their variable importance and threshold the voxel coefficient map so that less important voxels are removed from the final predictor. This final predictor can be used to obtain a ‘concreteness score’ for each brain image by calculating the dot product between the predictor and the image. We chose the number of PLS components to use and the level of thresholding based on the combination that gave the highest 24-fold leave-one-person-out cross validation prediction performance as measured by the area under the receiver operating characteristic curve (AUC).

**Figure 1.**
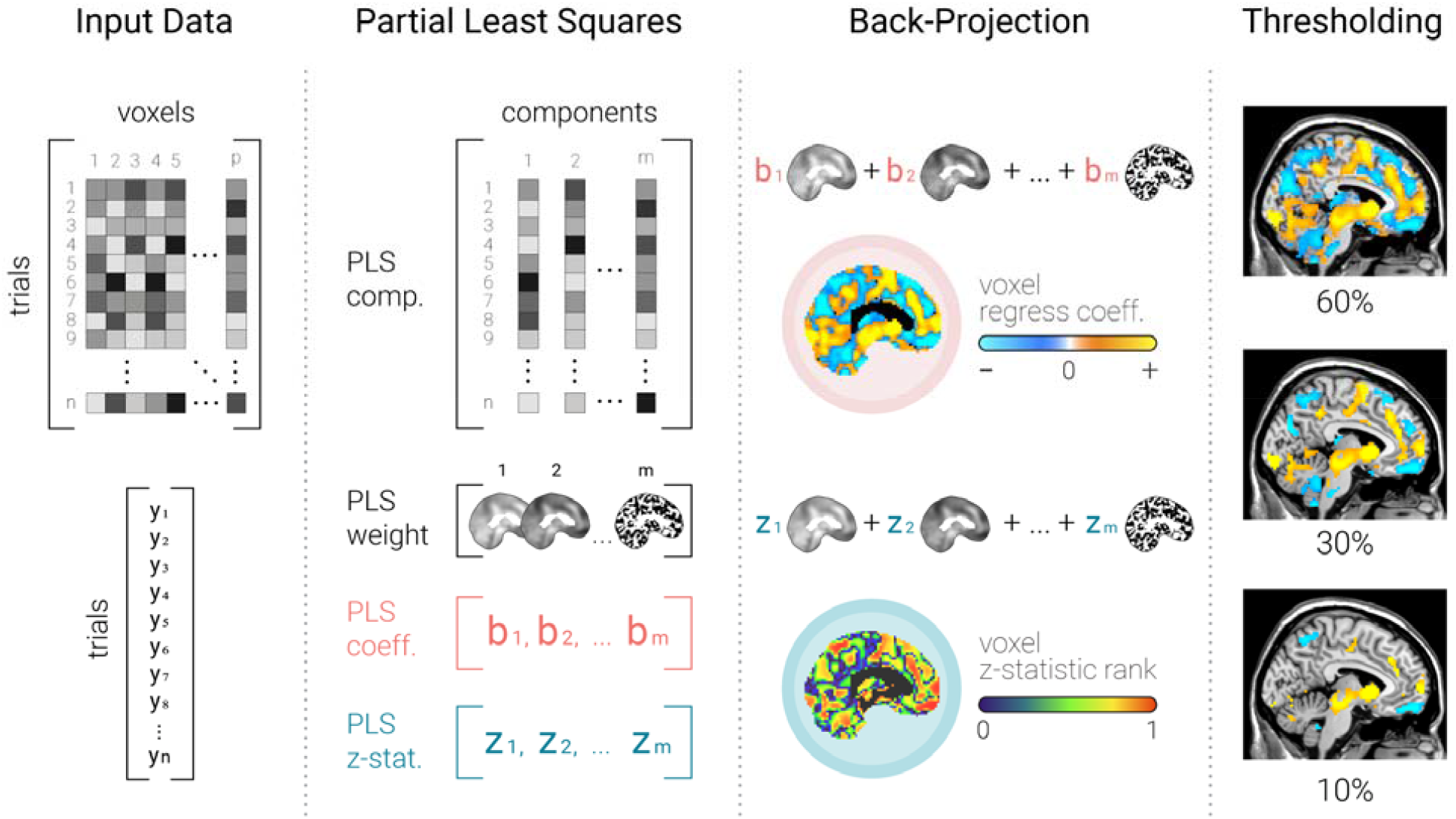
Thresholded Partial Least Square (T-PLS) approach to building a whole-brain predictor. Adapted with permission from (Lee et al., 2021). From left to right, the T-PLS method is outlined. The first step performs partial least squares on the brain image data (X) and the dependent variable (Y) in order to extract components that maximally explain the variance between X and Y. Each of these components are paired with weight maps that describe how each component is a weighted sum of the original voxels. They are also associated with regression coefficients and t-statistics (approx. z-stat) from regressing the dependent variable onto the components. These regression coefficients and z-stats are multiplied with their respective weight maps to yield regression coefficients and z-stats in the original voxel space. Using the voxel-level z-stats, the whole-brain predictor is thresholded by removing less important voxels (i.e., voxels with smaller absolute z-stats).

### Sensitivity and specificity analysis

To assess the accuracy of our whole-brain predictor of concreteness, we performed a nested 24-fold leave-one-person-out cross validation within the prospection dataset. We trained the predictor on data from 23 participants and tested on the one left-out person. Within the 23 training participants’ data, we employed an additional 23-fold leave-one-person-out cross validation to find the optimal number of components and thresholding level. After the best parameters were found, the T-PLS model was fitted using all 23 participants and used to predict the left out person’s data. Specifically, we tested whether the T-PLS model can accurately classify the high vs. low concreteness trial categories. Furthermore, we also tested if the T-PLS model predictions are correlated with the participants’ ratings of concreteness.

As we trained the T-PLS model on the condition labels for concreteness, and these were orthogonal by design to the condition labels for valence, we expected our whole brain predictor to be specific to concreteness and not valence. To assess the specificity of our whole brain predictor of concreteness, we also tested whether the T-PLS model could not accurately classify the positive vs. negative valence trial categories, and whether its predictions are not correlated with the participants’ ratings of valence.

### Concreteness and Delay Discounting

To calculate an expression score for the neural signature of concreteness during delay discounting, we calculated the dot product between the neural predictor of concreteness and the brain image of estimated activity for each trial. These scores were then correlated with the delay until the receipt of the delayed reward (in days), and the delayed amount (for Kable et al. 2017 only, since the delayed amount is constant in Cooper et al. 2013). The correlations were performed at the individual level, and each individual’s correlation coefficient was used as a summary statistic to test if there was a significant correlation at the group level.

## Results

We first developed a whole-brain neural predictor of the concreteness of prospective thought. We used an fMRI data set of 24 participants imagining possible future events that had been categorized *a priori* as high versus low in concreteness and positive versus negative in valence (Lee et al., 2020, **Fig 2A**). We used thresholded partial least squares (T-PLS, see **Fig. 1**) to develop a whole-brain classifier that discriminated events that were high versus low concreteness. We checked by cross-validation within the training dataset to ensure that our predictor of concreteness could accurately, out-of-sample, predict the concreteness but not the valence of imagined future events. As expected, the neural predictor of imagination concreteness successfully discriminated the trial categories of high versus low concreteness (mean prediction AUC = 64.76%, t-test against 50%, *t*(23) = 8.03, *p* < .0001), but not the trial categories of positive versus negative valence (mean prediction AUC = 51.16%, t-test against 50%, *t*(23) = 0.66, *p* = .51; **Fig. 2B**). We also further checked whether our predictions were also aligned with the participants’ ratings of the concreteness of imagined future events but not with their ratings of valence. Again, we found that our predictor was able to predict out-of-sample ratings of concreteness but unable to predict ratings of valence (**Fig. 2C**). Mean out-of-sample correlation between the neural prediction score and concreteness ratings was *r* = 0.16 (*t*(23) = 4.19, *p* = .0004), while the correlation between the neural prediction score and valence ratings was *r* = 0.0021 (*t*(23) = 0.063, *p* = .95).

**Figure 2.**
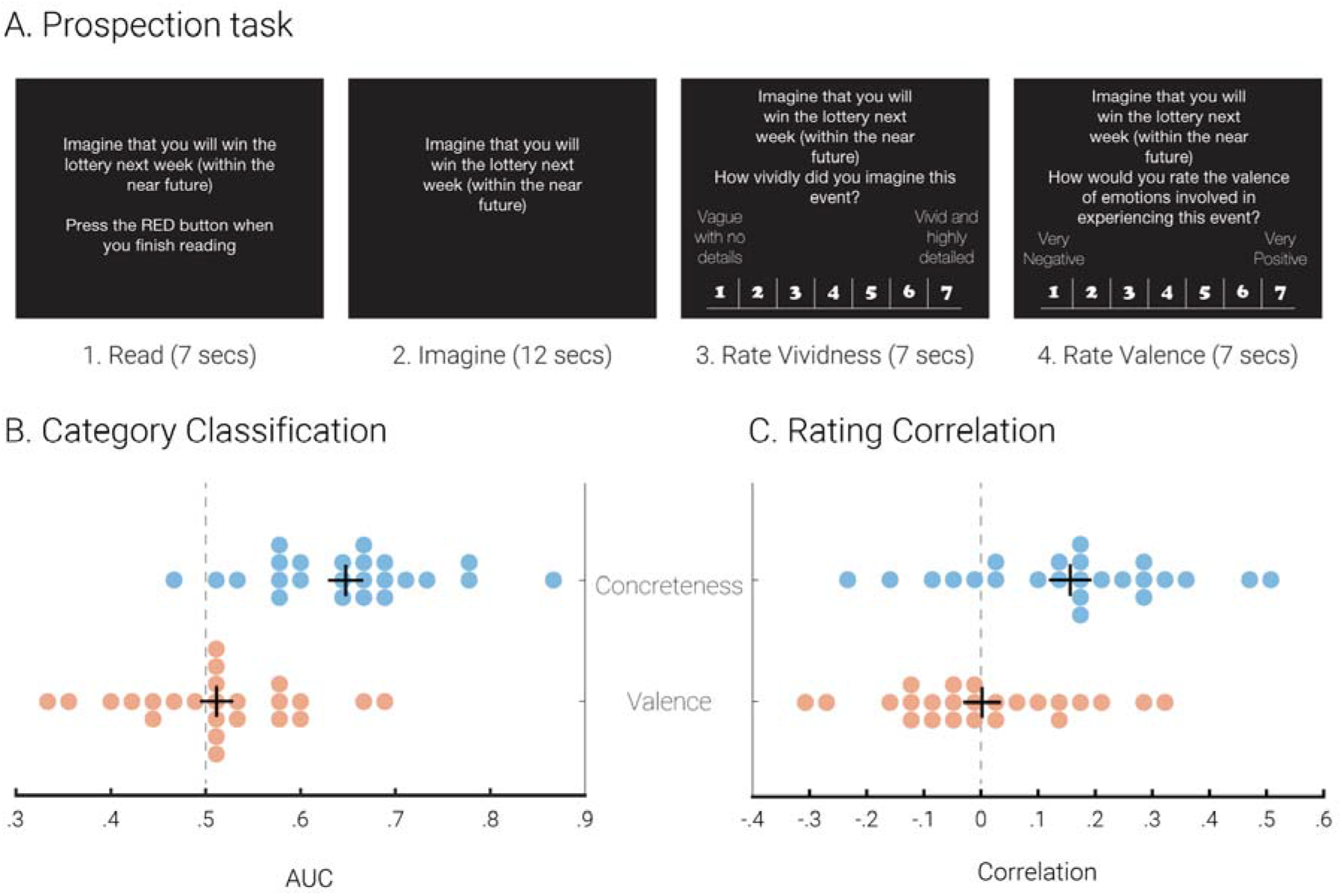
Out-of-sample prediction of concreteness and valence in the prospection dataset. Panel A shows the schematic of the prospection task from Lee et al. (2020). A whole-brain concreteness predictor is trained on 23 people’s data and used to predict the left-out person’s data. Panel B shows classification performance on *a priori* trial categories of concreteness (high versus low) and valence (positive versus negative) as measured by area under the receiver operating characteristic curve. Panel C shows correlation with concreteness and valence ratings provided by participants. Each dot represents one participant, the vertical bar represents the mean, and the horizontal bar represents the standard error of the mean.

The whole-brain prediction map of concreteness involved various regions of the brain, mostly in a bilateral fashion (**Fig. 3 & Table 1**). Positive coefficients (predictive of higher concreteness) were found in bilateral hippocampus, bilateral central orbitofrontal cortex (OFC), bilateral middle occipital gyri, right dorsolateral prefrontal cortex (dlPFC), and right inferior temporal gyrus. Negative coefficients (predictive of lower concreteness) were found in bilateral temporal poles, bilateral temporoparietal junction (TPJ), precuneus, and right cerebellum.

**Figure 3.**
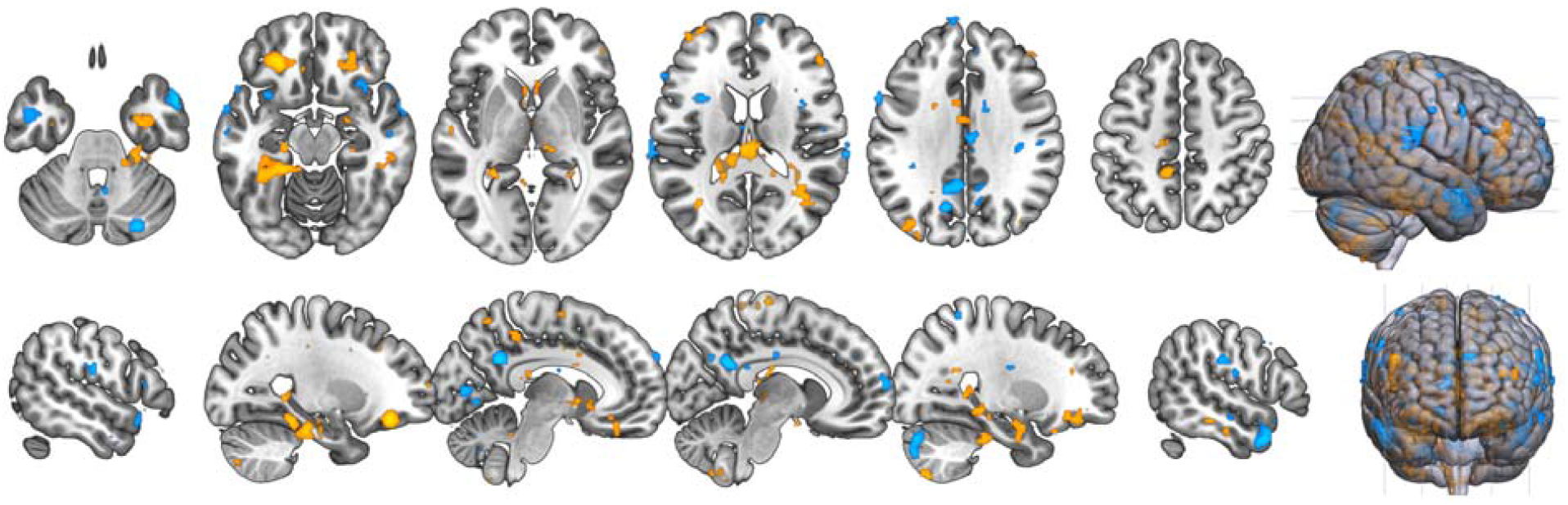
Whole-brain predictor of the concreteness of imagined future events. The warm colors indicate positive coefficients and cool colors indicate negative coefficients. Notable regions with positive coefficients include bilateral central OFC and bilateral hippocampus, and and with negative coefficients include precuneus, TPJ, and bilateral temporal pole.

**Table 1.** Clusters in the whole-brain predictor of imagination concreteness. Reported are clusters of voxels that have non-zero coefficients in the final predictor of the concreteness of imagined future events, grouped by the sign of the coefficients and ordered by cluster size. From left to right, the region names, cluster size in voxels, and peak MNI coordinates are provided. Clusters that are 50 voxels or smaller are not reported.

| Description | Size (number of voxels) | X | Y | Z |
| --- | --- | --- | --- | --- |
| *Positive* |  |  |  |  |
| Bilateral Hippocampus | 583 voxels (left) | -35 | -42 | -12 |
|  | 110 voxels (right) | 26 | -4 | -18 |
| Bilateral Central Orbitofrontal Cortex | 182 voxels (left) | -25 | 35 | -18 |
|  | 110 voxels (right) | 20 | 33 | -10 |
| Bilateral Middle Occipital Gyri | 109 voxels (left) | -35 | -78 | 37 |
|  | 230 voxels (right) | 40 | -60 | 17 |
| Right Dorsolateral Prefrontal Cortex<br>(right middle frontal gyrus) | 117 voxels | -39 | 37 | 11 |
| Right Inferior Temporal Gyrus | 198 voxels | 52 | -32 | -16 |
| *Negative* |  |  |  |  |
| Bilateral Temporal Pole | 64 voxels (left) | -47 | 0 | -28 |
|  | 167 voxels (right) | 58 | 9 | -28 |
| Bilateral Temporoparietal Junction | 98 voxels (left) | -69 | -30 | 15 |
|  | 55 voxels (right) | 70 | -30 | 17 |
| Bilateral Precuneus | 171 voxels (left) | -7 | -52 | 29 |
|  | 52 voxels (right) | -13 | -66 | 35 |
| Right Cerebellum | 118 voxels | 22 | -82 | -42 |

We next applied this whole-brain predictor of concreteness in two separate delay discounting tasks, in order to test whether the neural signature of concrete future thinking was higher when considering sooner rewards and lower when considering later rewards. In both datasets we found that neural concreteness scores were negatively correlated with the delay until the receipt of the reward, such that farther delays were associated with lower concreteness scores. Firstly, in an intertemporal bidding task, participants (n = 39) were presented with a fixed monetary outcome of $75 at different delays and asked to report the immediate amount they would feel to be equivalent to the delayed outcome. For each trial, we calculated neural concreteness scores by applying the whole brain predictor developed above to the activity for that trial. We found that the trial-by-trial neural concreteness scores were correlated negatively with delay (**Fig. 4**; mean *r* = −0.069, *t*(38) = −3.48, *p* = .0013), such that shorter delays (i.e., more proximal future) were associated with higher concreteness scores, and longer delays (i.e., more distant future) with lower concreteness scores.

**Figure 4.**
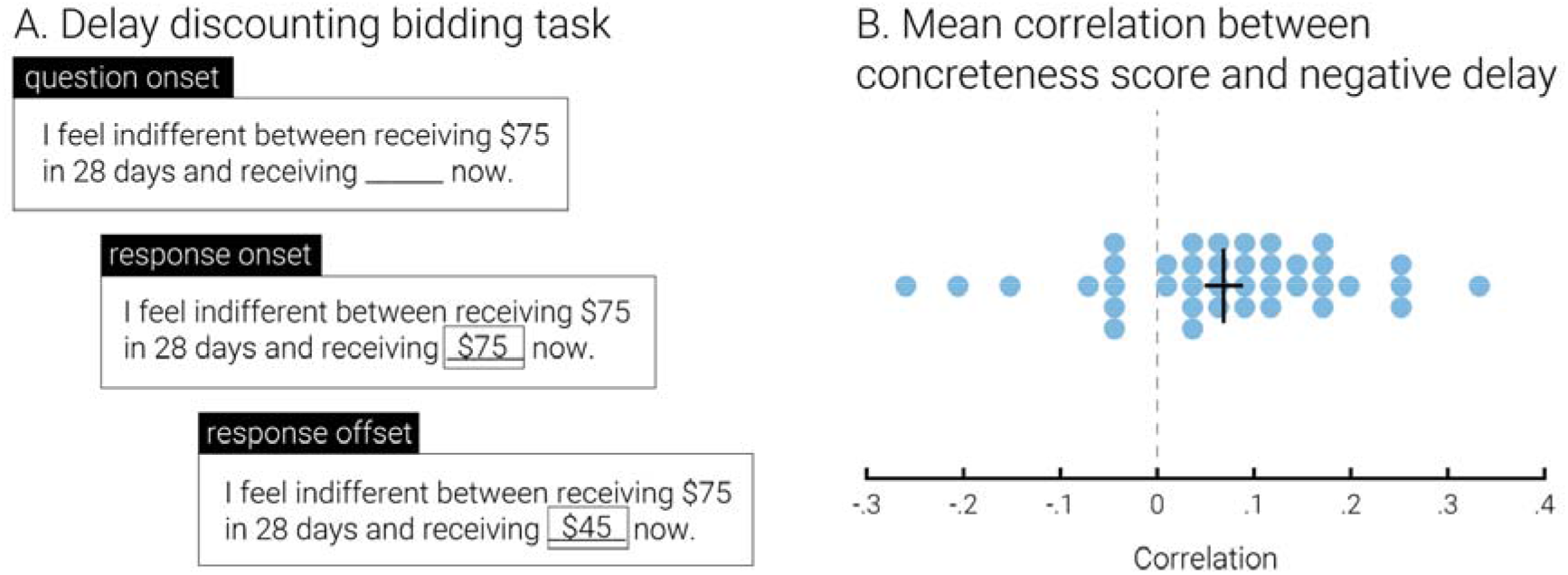
Out-of-sample prediction of delay in an intertemporal bidding task. Panel A shows the bidding task structure from Cooper et al. (2013). Participants are first shown the delayed amount of $75 (fixed) and a variable delay and are asked to bid their immediate equivalent. Panel B shows the per-person correlation between trial-by-trial delay (sign-flipped) and concreteness prediction scores from the whole-brain predictor. The vertical bar represents the mean, and the horizontal bar represents the standard error of the mean (n = 39).

We replicated this finding in a second dataset in which participants made discrete binary choices between immediate and delayed rewards. In this choice task from Kable et al. (2017), participants (n = 166) made choices between a fixed immediate reward of $20 and a future reward that varied in amount ($21 ∼ $85) and delay (20 ∼ 180 days) across trials. Again, we found that the trial-by-trial neural concreteness scores were correlated negatively with delay (**Fig. 5**; mean *r* = −0.050, *t*(165) = −6.09, bonferroni *p* < .0001), such that shorter delays were associated with higher concreteness scores. Furthermore, this association was specific to the delay to reward. The neural concreteness scores were not significantly correlated with the delayed amount (mean *r* = 0.016, *t*(165) = 2.12, bonferroni *p* = .11) and concreteness was more strongly associated with delay than amount (paired *t*-test, *t*(165) = 3.10, bonferroni *p* = .007).

**Figure 5.**
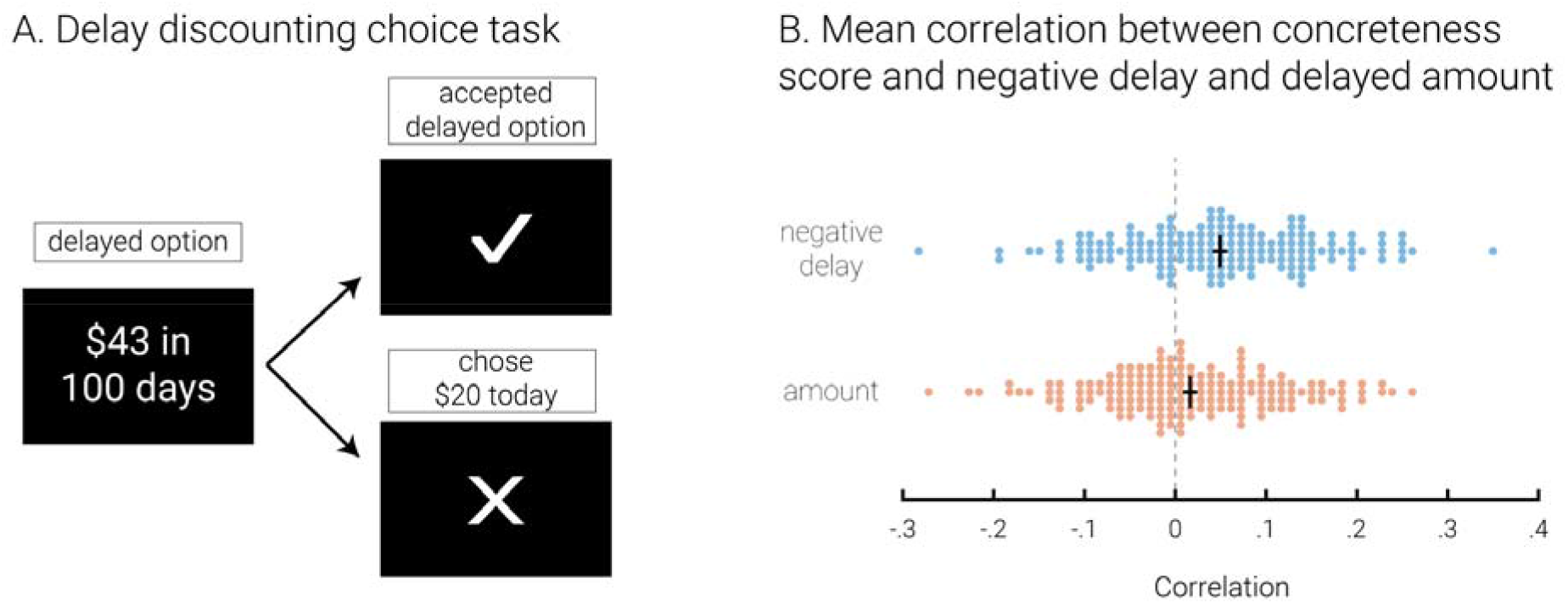
Out-of-sample prediction of delay in an intertemporal choice task. Panel A shows the choice task structure from Kable et al. (2017). Participants are shown the delayed reward and are asked to either accept it or to reject it for $20 immediately. Panel B shows the per-person correlation between trial-by-trial delay (sign-flipped) and concreteness prediction scores in comparison to that between trial-by-trial amount and concreteness prediction scores (delay has been sign-flipped to facilitate this comparison). The vertical bar represents the mean, and the horizontal bar represents the standard error of the mean (n = 166).

## Discussion

Multiple theories in psychology have suggested that delayed outcomes are discounted in value relative to immediate outcomes in part because more temporally distant options are perceived as less concrete and tangible than more temporally proximal ones (Liberman & Trope, 2014; Rick & Loewenstein, 2008; Trope & Liberman, 2010). These theories have been supported by a range of various behavioral experiments (Bischoff & Hansen, 2016; Kelley & Schmeichel, 2015; Liberman & Trope, 2014; Malkoc et al., 2010; Mischel & Baker, 1975; Yi, Stuppy-Sullivan, Pickover, & Landes, 2017). Here we add converging neuroscientific evidence to these theories.

We used fMRI data during an imagination task to create a whole-brain, multivariate predictor specific to the concreteness of prospective thought, independent of the valence of prospective thought. Then we show, in two separate delay discounting datasets with markedly different task structure (one bidding task, one choice task), that the neural signature of concreteness is modulated by the temporal distance to the delayed option under consideration. That is, the pattern of neural activity that predicts more concrete prospective thinking is stronger for more temporally proximal outcomes and weaker for more temporally distal ones. The neural signature of concreteness was also more strongly modulated by the delay to reward than by the magnitude of reward. These results show that, while people are making intertemporal decisions, an online, unobtrusive neural index of concrete thinking declines as the outcomes considered are delayed farther into the future.

Our results complement previous tests of construal level theory using fMRI. These studies have shown that neural activity associated with imagining near events, compared to distant events, overlaps with neural activity engaged by other forms of psychological proximity or by low-versus high-level construal (Stillman et al., 2017; Tamir & Mitchell, 2011). Here we make two advances over these previous results. First, we distinguish between neural activity due to the concreteness, versus the valence, of prospective thought. This is critical as several previous studies have found the strongest increases in activity for sooner, compared to later, events in the mPFC and PCC (Mitchell et al., 2011; Tamir & Mitchell, 2011), two regions that we have previously shown are associated with the valence of prospective thought (Lee et al., 2020). Second, we show that a neural index of concreteness is modulated by the delay to the outcome *during intertemporal decision-making*. This links reduced concreteness directly to the discounting of future rewards, a process known to be associated with many important life outcomes (Alessi & Petry, 2003; Anderson & Mellor, 2008; Brañas-Garza et al., 2007; Kirby et al., 1999; Krain et al., 2008; Lejuez et al., 2005, 2003; Schepis et al., 2011; Shamosh & Gray, 2008).

The whole-brain prediction map for concreteness is remarkably consistent with findings from other lines of research. Several previous studies have argued that the orbitofrontal cortex represents the features of potential outcomes during decision making (Burke, Franz, Miller, & Schoenbaum, 2008; Howard, Gottfried, Tobler, & Kahnt, 2015; Takahashi et al., 2013), and that interactions with the hippocampus may be critical for generating these representations from memory (for review, see Shohamy & Daw, 2015). Furthermore, there is evidence that these regions play a role in valuing delayed rewards. Lesions to the OFC caused increased impatience (Sellitto, Ciaramelli, & Di Pellegrino, 2010), and reduced grey matter thickness in both the OFC (Pehlivanova et al., 2018) and the medial temporal lobe (Lempert et al., 2020; Owens et al., 2017) is associated with increased discounting. Correspondingly, we would expect that modulating activity in these regions as people consider future outcomes would alter the concreteness with which those outcomes are imagined and the degree to which those outcomes are discounted.

To obtain the current results, we applied a novel adaptation of partial least squares optimized to construct interpretable whole-brain predictors with minimal computation time (Lee et al., 2021). Though many different methods for constructing whole brain predictors have been proposed (Grosenick, Greer, & Knutson, 2008; Kragel & LaBar, 2014; Smith, Douglas Bernheim, Camerer, & Rangel, 2014; Wager et al., 2013), none have yet achieved widespread use in the field. Here we illustrate what we think is the most promising and exciting potential use of such predictors: decoding mental states online in order to test psychological hypotheses.

